# Targeting G protein-coupled receptor-17 (GPR17) upregulation in paediatric diffuse midbrain gliomas leads to altered phenotype and susceptibility to therapies

**DOI:** 10.1101/2020.11.17.386706

**Authors:** Katie F. Loveson, Helen L. Fillmore

## Abstract

Paediatric diffuse midline glioma (pDMG) also known as Diffuse intrinsic pontine gliomas (DIPG) is an incurable, aggressive childhood brain malignancy, that arises in a region- and age-specific nature. The underlying pathophysiology suggests dysregulation of postnatal neurodevelopmental processes causing aborted cell differentiation. The cell of origin is unclear, but data suggests an oligodendrocytic lineage (OPC), supported by the over-expression of transcription factors such as Olig1 and Olig2 in 80% of DIPG cases.

In-depth bioinformatics and principal component analyses (PCA) of genes involved in brain development and pDMG support reports of OPC gene dysregulation and led to the identification of the G-protein coupled receptor 17 (GPR17) and its association with pDMG. GPR17 mRNA and protein expression was confirmed in all pDMG cell lines tested. Using a well-characterised agonist (MDL 299,51) and antagonist (HAMI3379) to modulate GPR17 function in pDMG cell lines resulted in phenotypic and genomic changes as well as in cell growth and migration. HAMI3379, a GPR17 specific antagonist resulted in a significant reduction in GPR17 mRNA and protein expression (p<0.006) and a significant reduction in migration (p<0.0025). When pDMG cells were pre-treatment with HAMI3379 in combination with known cytotoxic agents (Bleomycin, a radiation mimic, Panobinostat or Vincristine), there was a decrease in cell viability compare to cytotoxic agent alone. There are no current effective therapies for pDMG patients and the ability of blocking GPR17 function to enhance sensitivity to standard therapies is appealing and warrants further investigation.

## Introduction

Brain tumours are the leading cause of cancer-related death in children and are the second most common malignancy in children, after leukaemia (1–4). Diffuse intrinsic pontine gliomas (DIPG), recently reclassified as paediatric diffuse midline glioma (pDMG – H3 K27M mutant), represent approximately 10% of childhood brain cancer, with medulloblastoma being the most common (20%) (5–7).

pDMG H3 K27M mutant, is a rare cancer, with a yearly incidence of 2.32 per 1,000,000 people aged 0-20 years (7). These tumours primarily affect young children with peak incidence at 6 years of age and have the highest mortality of all childhood solid tumours (8). The pontine (pons) region of the brainstem where these tumours arise, increases 6-fold in size from birth to 5 years of age, with continued growth, albeit slower throughout childhood (9). This is likely due to increased myelination and the development of neural circuits through learned behaviour (8–11). Growth of pDMGs occur in a discrete spatial and temporal pattern and coincides with periods of developmental myelination (12,13).

We used a bioinformatic approach to examine genes associated with myelination in brain development followed by an assessment of their expression in publically available pDMG patient platforms. We identified the G-protein coupled receptor 17 (GPR17) to be dysregulated in pDMG, alongside known genes PDGFRA, CSPG4 and OLIG2. GPR17 is a rhodopsin-like orphan GPCR (unknown ligand) and has the typical features of the GPCR superfamily of seven transmembrane domains (TM1-TM7), eight amphipathic helices, an extracellular n-terminal domain and an intracellular C-terminus (14–17). GPR17 is predominantly expressed in the Central Nervous System (CNS) and has been implicated in a number of normal physiological and pathological processes, such as oligodendrocyte differentiation, spinal cord injury, brain injury and tissues that undergo ischemic damage such as the kidney and heart (14,16,18–20).

GPR17 is expressed in OPCs in a controlled manner and is seen as an intrinsic cell timer of differentiation as well as a sensor for injury (17,19,21,22) Studies suggest that GRP17 is vital to the transition between immature and myelinating oligodendrocytes via, ID2/4 mediated negative regulation (22–26). Expression of GPR17 is regulated through the oligodendrocyte pathway; with the mechanism still not entirely clear (Figure 1A). Single cell RNA-sequence studies of the mouse and human midbrain have identified GPR17 to be almost exclusively expressed by OPCs (Figure 1B) (23). *In vitro* models report that GPR17 overexpression in transgenic mice results in severe myelination defects, whereas GPR17 null mice exhibit accelerated differentiation and early onset of myelination. The development of therapies targeting GPR17 has been largely focused on neurodegenerative disorders and brain injury (7,8,23) and can therefore be used to investigate the potential role of GPR17 in pDMG pathology.

**Figure 1.**
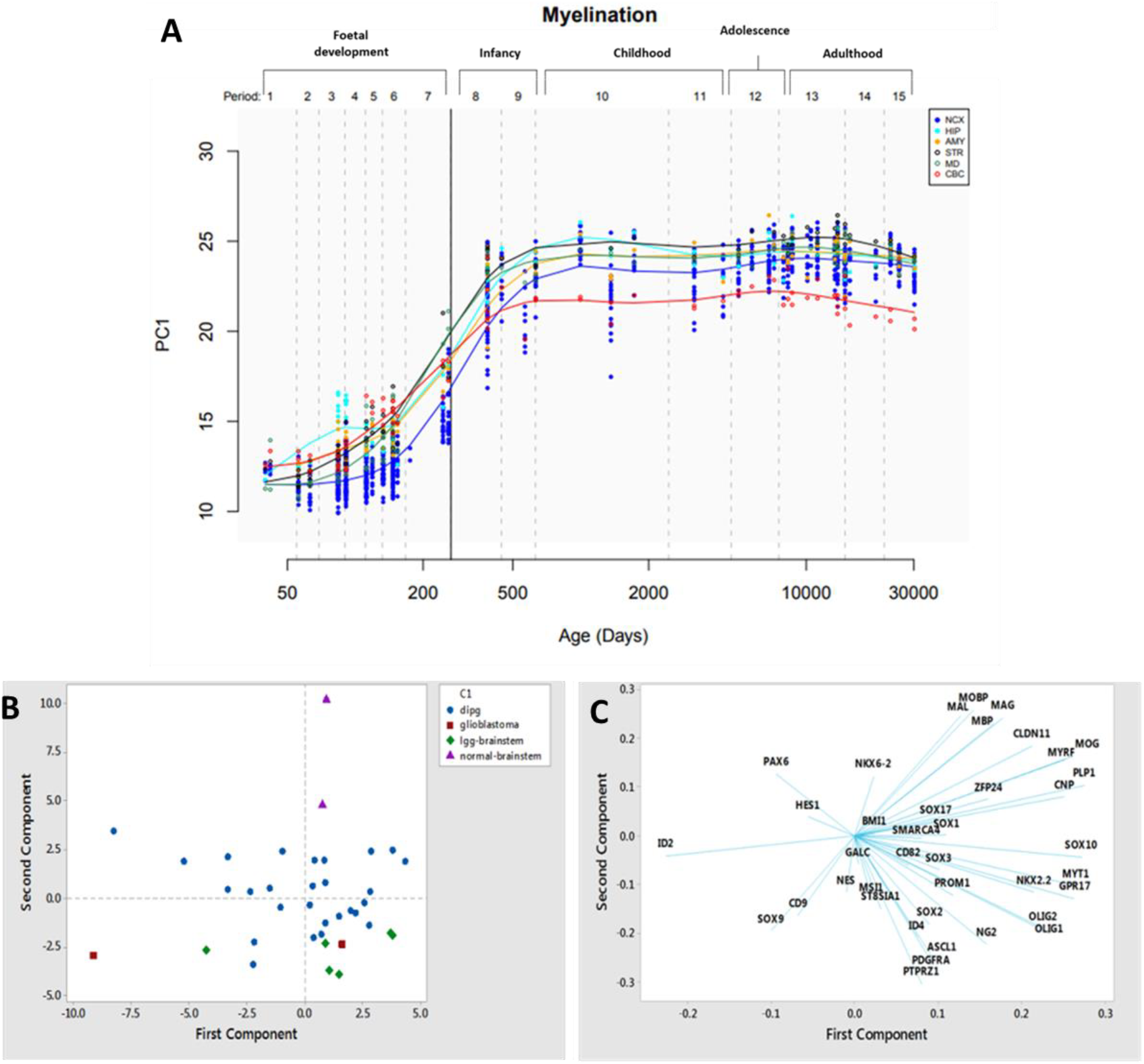
Developmental trajectory of genes associated with myelination demonstrating specific spatiotemporal gene expression profiles from foetal development to adulthood. A) PCA of the expression of grouped genes (MBP, PLP1, MOG, C11orf9, MAG) in cerebellar cortex (CBC), mediodorsal nucleus of the thalamus (MD), striatum (STR), amygdala (AMY), hippocampus (HIP) and neocortex (NCX). Data obtained from (Kang et al., 2011). B) PCA Analysis of stage-specific Oligodendrocyte markers across different histologically grades. Score plot of stage specific OL lineage markers. High grade tumours cluster away from normal brainstem and low-grade brain tumours. C) Loading plot of the first two components, genes shown to be higher in DIPG cluster in the bottom right quadrant and genes associated with normal brainstem, in the top right. Genes associated with OPC stage, cluster together.

Herein, we used a two-pronged approach using bioinformatic analyses of publicly available database platforms to provide evidence of dysregulation of GPR17 in pDMG and inform our *in vitro* studies. We demonstrate that modulating its expression using a well-characterised agonist (MDL 299,51) and antagonist (HAMI3379) led to changes in pDMG cell line growth and migration. pDMG cell lines treated with HAMI3379 were more responsive in terms of cell viability to Bleomycin (<u>radiation</u> mimic), Panobinostat or Vincristine. Overall, our data suggest that blocking GPR17 caused pDMG cells to become more sensitive to standard therapies.

## Results

### Genes involved in the Oligodendrocyte Lineage pathway are dysregulated in DIPG

Several genes involved in myelination (MBP, PLP1, MOG, C11orf9 and MAG) were examined in a spatiotemporal manner using the Human Brain Transcriptome Atlas (24). There is an increase in the developmental trajectory from the last third of gestation to 3 years of age, where it remains constant into adulthood (Figure 1A).

The cells responsible for myelination in the central nervous system are specialised glia cells, oligodendrocytes. Using Principal Component Analysis (PCA), we analysed expression of genes involved in the oligodendrocyte lineage pathway in an available pDMG dataset (25). As shown in Figure 1B, the brain tumour patients cluster away from normal controls, in particular pDMG and low-grade brain tumours. The loading plot from the PCA was examined to understand the loading of individual genes on the principal components (Figure 1C). Here we identified GPR17 to cluster with genes known to be upregulated in pDMG: *PDGFRA, OLIG1* and *OLIG2* which are key in the oligodendrocyte pathway. These genes are up-regulated in pDMG patients compared to normal controls suggesting that pDMG have some commonality with OPCs (26). Clustering of all stage-specific oligodendrocyte markers are shown in Supplemental Figure S1.

### GPR17 is dysregulated in DIPG

Initial bioinformatic analyses indicated an increase in GPR17 mRNA expression in pDMG (n=27) compared with normal brain (all regions, n=172) and normal brainstem (n=2) (Figure 2A). Due to the limitation of access to normal paediatric brainstem tissue we included the ‘all brain regions’ category for comparison. Due to the importance of the histone mutation K27M (H3.1 and H3.3) in pDMG, we also examined the association of GPR17 mRNA expression with these mutations and found that there was a significant increase in GPR17 expression in all K27M (H3.1 and H3.3) subtypes compared to control (Figure 2B). Data mining of single cell RNA-Seq studies of the mouse and human midbrain have identified GPR17 to be almost exclusively expressed by OPCs (Figure 2C) (23).

**Figure 2.**
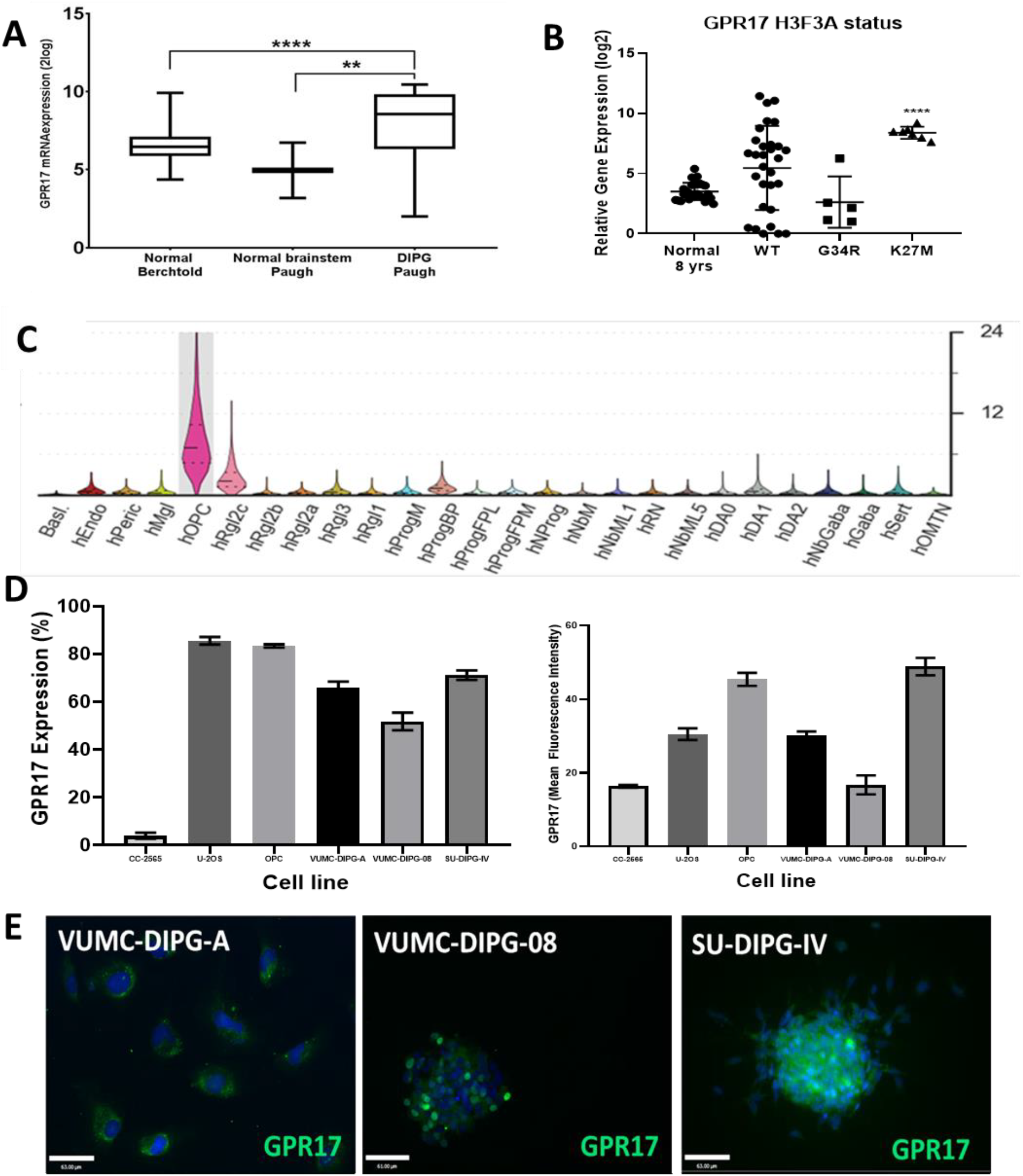
Bioinformatic analysis and in vitro characterisation of GPR17. A) GPR17 is up-regulated in pDMG patients compared to normal brain and brainstem **** p<0.0001 ** p 0.0037 (GSE26576) (Paugh et al., 2011). B) GPR17 is overexpressed in K27M tumours in comparison to normal age matched brain. Statistical analysis performed using GraphPad Prism, One-way ANOVA, followed by Dunnett’s multiple comparison test **** p<0.0001 ** 0.0052 (Datasets: Pfister, GSE36245 and Berchtold) (Sturm et al., 2012). C) Violin plot of single-cell RNA-sequence studies show GPR17 to be predominately to be expressed by OPCs (23). D) Flow cytometry analysis of GPR17 in non-neoplastic and DIPG cell lines. U-2OS used as a positive control. A) Percentage of GPR17 expression. C) Mean Fluorescence Intensity of GPR17 of per cell. E) Immunostaining of OPC and DIPG cells shows expression of GPR17 Fluorescence images of DIPG cell lines, GPR17 (Green) and nuclei stained blue. Images taken at 20x magnification, and 63μm (DIPG).

We further characterised GPR17 expression in primary pDMG cell lines and non-neoplastic controls at the protein level using flow cytometry and immunocytochemisty. The mean percentage of cells GPR17 positive were found to be similar in the three pDMG cell lines tested, ranging between 51.82 and 71.17% (VUMC-DIPG-A, 66.06%, VUMC-DIPG-08, 51.82%, and SU-DIPG-IV, 71-17%) compared to GPR17 positive expression in positive control cells, U-2OS, 85.57% and OPCs, 83.42%; and compared to less than 3.97% in the CC-2565 foetal astrocyte cell line (Figure 2D). The VUMC-DIPG-08 cell line showed the lowest, then VUMC-DIPG-A and SU-DIPG-IV having the highest percentage (Figure 2D). The mean fluorescence intensity (MFI) of the pDMG cell lines had a similar trend as the expression, with SU-DIPG-IV (48.86) having the highest MFI, followed by VUMC-DIPG-A (30.2) and VUMC-DIPG-08 (16.73) (Figure 2D). Protein expression was also confirmed in the pDMG cell lines using immunocytochemistry (Figure 2E).

### Treatment with HAMI3379 alters the cellular phenotype

To explore the potential function of GPR17 two experimental drugs were used for *in vitro* experiments. The small molecule agonist, MDL 299,51, highly selective for GPR17 and a GPR17 antagonist, initially developed as a CysLT2 antagonist to treat cardiovascular and inflammatory disorders were utilized in our investigations (12,27). The OPCs and VUMC-DIPG-A cell line treated with 10nM MDL 299,51 for 48 hours had a similar morphology to untreated controls (Figure 3B). The two cell lines that grow as suspension spheres, VUMC-DIPG-08 and SU-DIPG-IV had larger spheres, compared to control when treated with 10nM MDL 299,51 (Figure 3B). Treatment of the cell lines with 30μM HAMI3379 resulted in the alteration in cell morphology of all cell lines, including non-neoplastic controls. The cells had an increased number of processes (arrows) and a reduction in the number of spheres (VUMC-DIPG-08 and SU-DIPG-IV) (Figure 3B).

**Figure 3.**
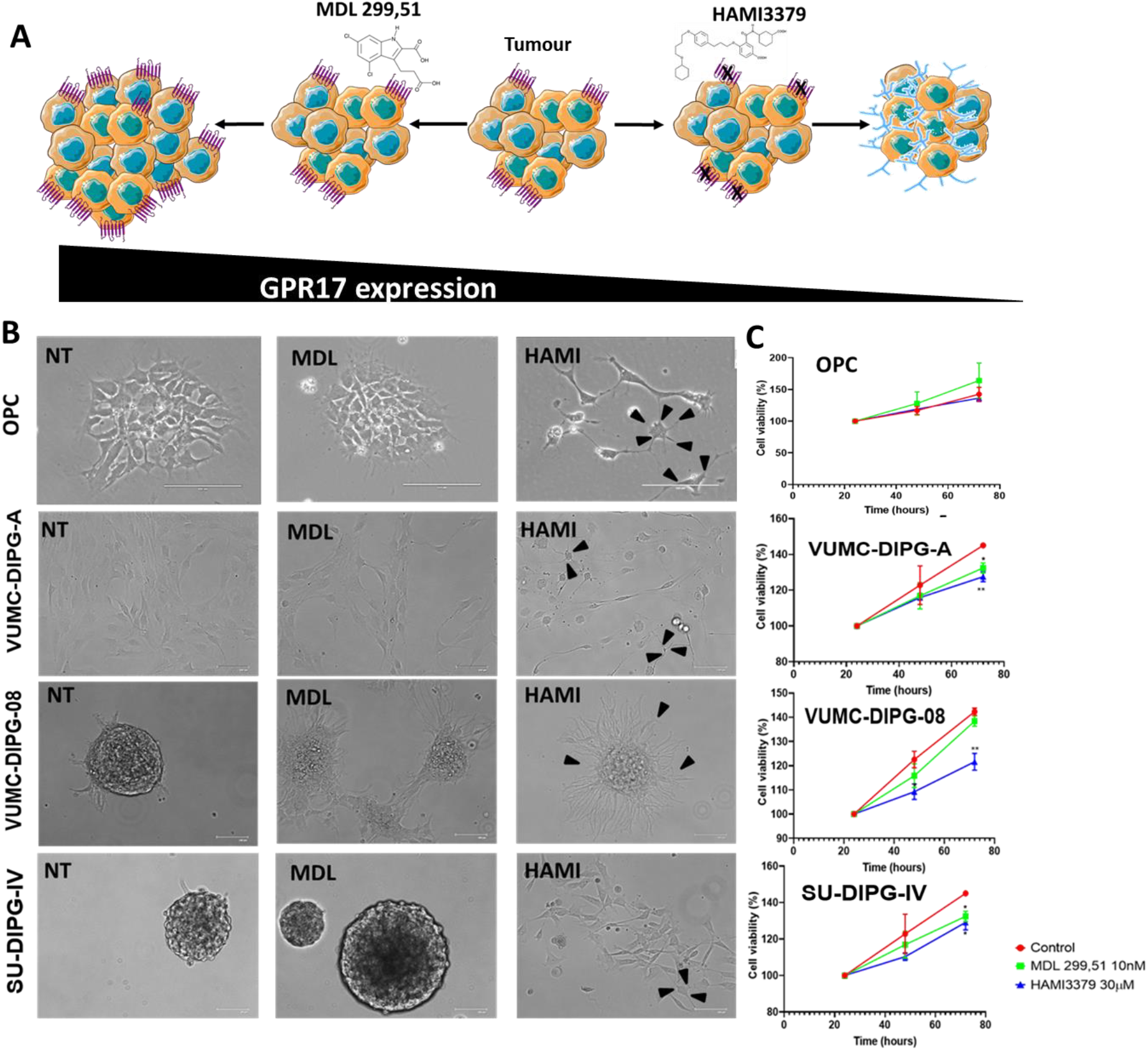
Treatment of cell lines with 10nM MDL 299,51 and 30μM HAMI3379 alter the cells morphology and viability. A) Schematic of the proposed mechanisms of MDL 299,51 and HAMI3379. MDL 299,51 activates GPR17 and keeps the cells in a proliferative state. Whereas HAMI3379, blocks GPR17 forcing the cells to differentiate. B) Representative images of cell lines treated with MDL 299, 51 and HAMI3379 for 48 hours. Brightfield mages taken using FLoid imaging system. Scale bar 100μm. C) Cell viability of cell lines when treated with GPR17 agonist (MDL 229,51, 10nM) or antagonist (HAMI3379 30μM) in normal growth conditions. MTS assay performed at 24, 48 and 72 hours after treatment and absorbance measured at 490nm. Measurements were blank corrected and normalised to 24-hour time point. Graphs show mean with SEM. Statistics were performed using GraphPad Prism, Two-way ANOVA with Dunnett’s multiple comparison test. Treatment with 30μM HAMI3379 caused a significant decrease in viability compared to control at 72 hours in all cell lines except OPC’s. *<0.03 **<0.007 ***<0.0008. Treatment with 10nM MDL 299,51 caused a significant decrease in viability compared to control at 72 hours in CC-2565, VUMC-DIPG-A and SU-DIPG-IV. *<0.03 ***<0.0008.

MDL 299,51 caused a significant decrease in cell viability in normal growth conditions of VUMC-DIPG-A and SU-DIPG-IV at 72 hours (p=0.03 and p=0.0008, respectively) compared to no treatment control (Figure 3C). Treatment with HAMI3379 caused a significant decrease in cell viability in all cell lines, except OPCs (p<0.05) compared to no treatment control (Figure 3C). Following analyses of these data, experiments were performed at the 48-hour time point as there was no significant effect on cell viability was observed.

### Treatment with MDL 299,51 and HAMI3379 alter the mRNA levels of GPR17 in pDMG cell lines

To determine whether the treatment of the cell lines with 10nM MDL 299,51 and 30μM HAMI3379 alters the mRNA expression of GPR17, RNA was extracted 48 hours after treatment with the compounds and examined using RT-qPCR. MDL 299,51 caused a significant up-regulation of GPR17 mRNA in VUMC-DIPG-A (p=0.0242) and SU-DIPG-IV (p=0.006) when compared with no treatment. Treatment with HAMI3379 resulted in a significant down regulation of GPR17 mRNA expression in VUMC-DIPG-A and VUMC-DIPG-08 (p=0.006) (Figure 4A).

**Figure 4.**
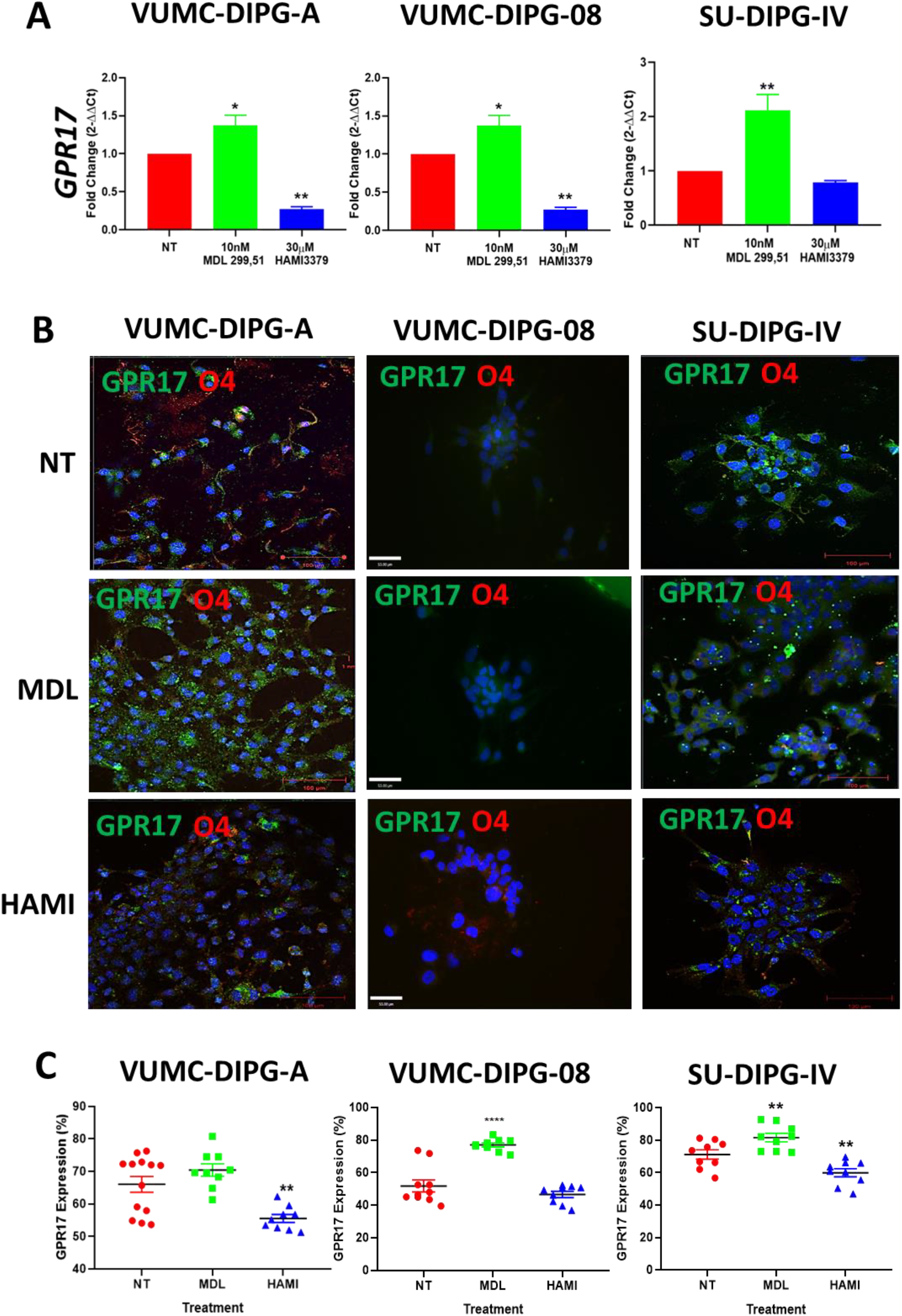
GPR17 expression is modulated using MDL 299,51 and HAMI3379. A) Relative GPR17 mRNA expression is increased in DIPG cell lines treated with 10nM MDL 299,51 and decreased when treated with 30μM HAMI3379 for 48 hours. Data were analysed by the 2 −ΔΔCt method using GAPDH as a reference gene. Bars show mean with SEM n=3. Statistics performed using GraphPad Prism. One-way ANOVA with Dunnett’s multiple comparison test. ** p<0.006 and *p =0.0242. B) Immunostaining of DIPG cell lines when treated with GPR17 agonist (MDL 229,51, 10nM) or antagonist (HAMI3379 30μM) for 48 hours. Confocal images of DIPG cell lines, GPR17 (Green), O4 (Red) and nuclei stained blue. Images taken at 20x magnification, scale bar 100μm. Localisation of GPR17 appears to be cytoplasmic in all treatments, with an increase in the MDL 299,51 treated cells. O4 staining is increased with treatment of HAMI3379. C) Flow cytometry analysis of percentage overall GPR17 expression in DIPG cell lines when treated with GPR17 agonist (MDL 229,51, 10nM) or antagonist (HAMI3379 30μM) for 48 hours. Graphs show mean with SEM. Statistics were performed using GraphPad Prism, One-way ANOVA with Dunnett’s multiple comparison test by comparing treated cells with untreated control. ** <0.006 **** <0.0001.

### pDMG cells treated with HAMI3379 leads to an increase in O4, a marker of Oligodendrocyte differentiation

When the DIPG cell lines are treated with HAMI3379, there was an altered cell morphology (Figure 4B), more like that of a pre-myelinating/mature oligodendrocyte like cell. Therefore, ICC was conducted ICC using Oligodendrocyte Marker O4 (O4) on cells that had been treated with either 10nM MDL 299,51 or 30μM HAMI3379 for 48 hours. Figure 4B shows an increase in O4 staining in the HAMI3379 VUMC-DIPG-A cells, whereas treatment with MDL 299,51 caused an increase in GPR17 staining. The changes in O4 expression in VUMC-DIPG-08 cells were much more subtle, but still increased compared to no treatment control (Figure 4B). The SU-DIPG-IV cells, like the other two DIPG cell lines had an increase in O4 expression when treated with HAMI3379. Interestingly the expression of GPR17 appears to be more consistent across the cells in MDL 299,51 treated compared to no treatment and HAMI3379, where it appears more punctate (Figure 4B).

### GPR17 protein levels are decreased upon treatment with HAMI3379

To determine the effect of compounds on GPR17 protein expression we used ICC and flow cytometry to access changes in expression. Treatment with MDL 299,51 caused a significant increase in percentage GPR17 protein expression in VUMC-DIPG-08 and SU-DIPG-IV (p<0.02) and a significant decrease in VUMC-DIPG-A and SU-DIPG-IV (p<0.02) (Figure 4C). The mean fluorescence intensity or number of antigens per cell was significantly increased in all cell lines when treated with MDL 299,51 (p<0.02) but only significantly decreased in SU-DIPG-IV (p<0.02) (Figure 4C).

### HAMI3379 reduces pDMG cell distance travelled

Due to the diffuse nature of pDMGs and to determine the functional effect of MDL 299,51 and HAMI3379, we next examined the motility of the cell types by tracking the distance travelled of single cells. Treatment with MDL 299,51 caused a significantly increase in average distance travelled in the OPCs and VUMC-DIPG-08 (p=0.0025 and p=0.0001, respectively, Figure 5). MDL 299,51 also caused a significant increase in mean velocity in the OPCs (No treatment; 2.35 μm/second compared to 2.91 μm/second) and VUMC-DIPG-08 (No treatment, 2.53 μm/second compared to 3.06 μm/second p<0.0025), whereas treatment with HAMI3379 caused a significant decrease in average distance travelled in VUMC-DIPG-08 and SU-DIPG-IV (p <0.0025, Figure 5).

**Figure 5.**
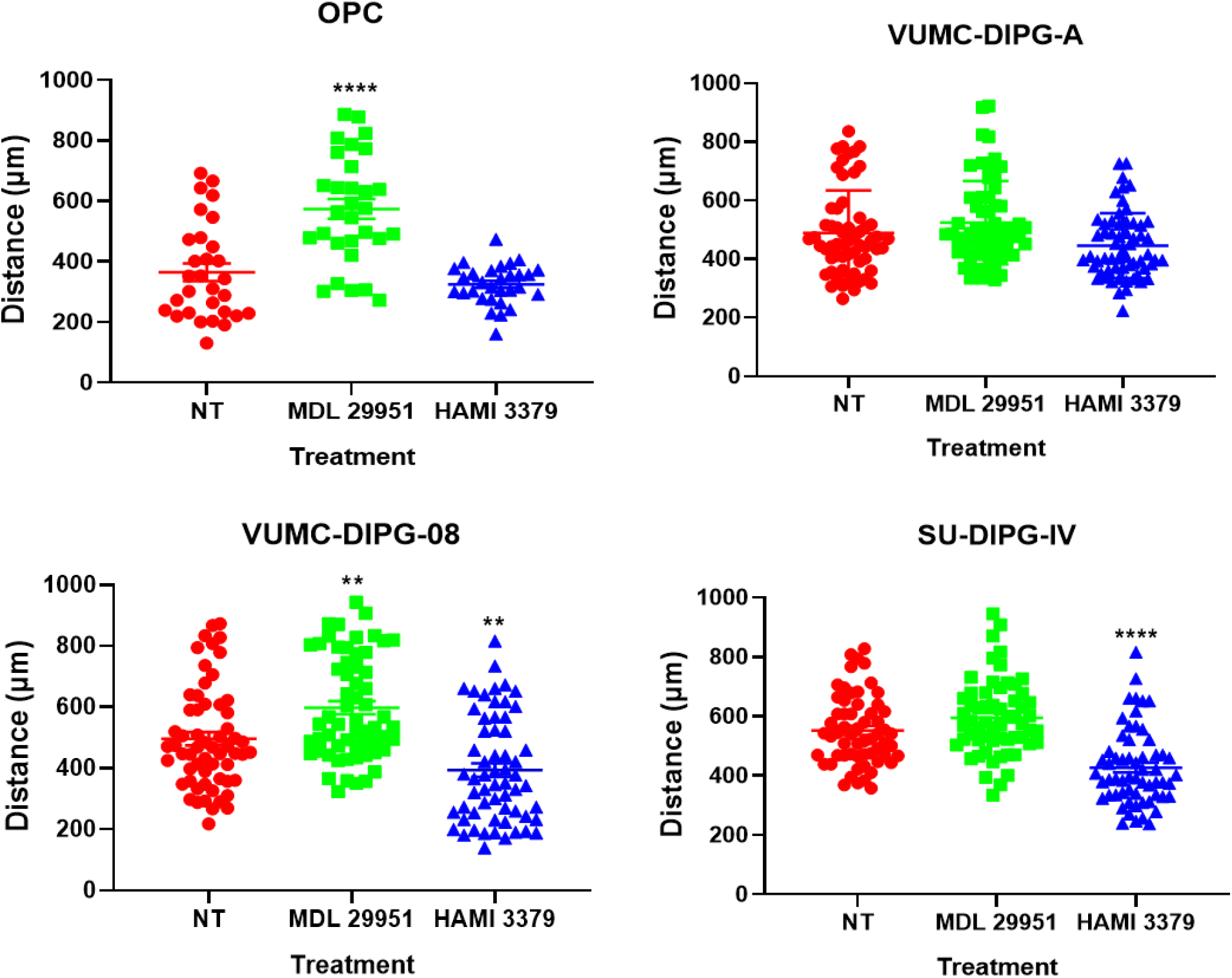
Average distance (μm) travelled by cell lines. Treatment with 30μM HAMI3379 caused a significant decrease in distance travelled compared to no treatment control for VUMC-DIPG-08 and SU-DIPG-IV. Whereas treatment with 10nM MDL 299,51 caused a significant increase in distance travelled compared to control in the OPCs and VUMC-DIPG-08. ** <0.0025 ****<0.0001.

### HAMI3379, a GPR17 specific antagonist causes pDMG cell lines to be more sensitive to existing therapies

When pDMG cells were treated with HAMI3379, GPR17 expression decreased and there was an observed change in cell morphology, as well as a reduction in self renewal capacity (Figure S2) and motility. To test our hypothesis that modulation of GPR17 would make DIPG cells more susceptible to existing therapies (Figure 6A), DIPG cells were treated with 30μM HAMI3379 for 24 hours, then with the EC50 dose determined for each compound as described in Supplementary Table 1. Prior treatment with HAMI3379 caused a significance decrease in cell viability in the VUMC-DIPG-A cells in combination with Panobinostat or vincristine compared to cytotoxic alone (p<0.0003, Figure 6A). VUMC-DIPG-08 had a significant decrease in cell viability with prior HAMI3379 treatment followed by Bleomycin or vincristine compared to cytotoxic agent alone (p=0.038 and p=0.0007, respectively,Figure 6B). When SU-DIPG-IV cells were pre-treatment with HAMI3379, there was a significant decrease in cell viability compared to cytotoxic alone with all treatments (p<0.01 and p=0.0001, respectively Figure 6C).

**Figure 6.**
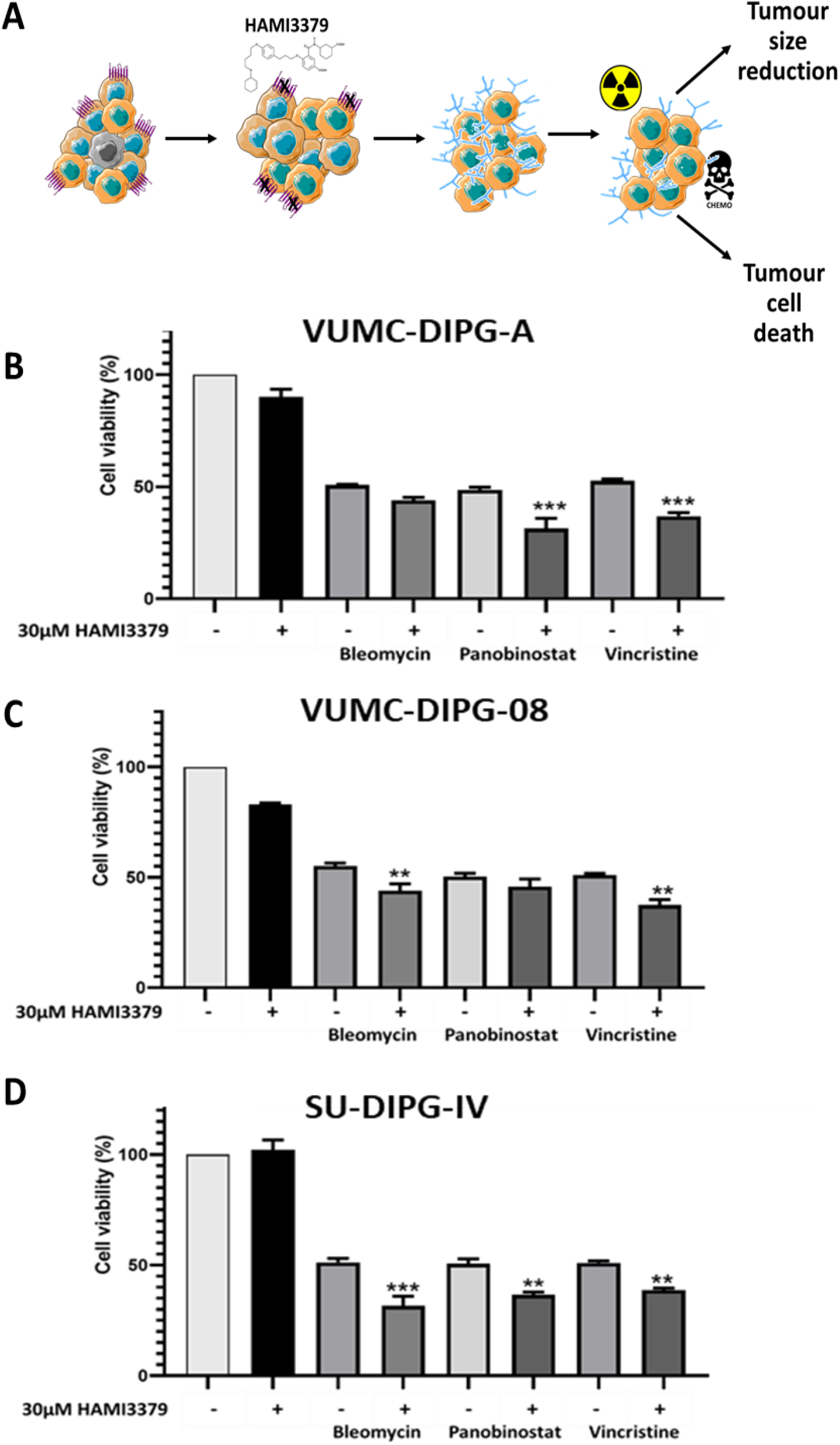
Cell viability of cell lines with prior treatment of 30μM HAMI3379 for 24 hours then and EC50s of Bleomycin, Panobinostat or Vincristine for 48 hours. A) Schematic of the proposed mechanism of HAMI3379. B) VUMC-DIPG-A show a significant decrease in cell viability with prior treatment of HAMI3379 followed by EC50’s of Panobinostat and Vincristine (***<0.0003). C) VUMC-DIPG-08 show a significant decrease in cell viability with prior treatment of HAMI3379 followed by EC50’s of Bleomycin and Vincristine (** p=0.038 and ***p=0.0007). D) SU-DIPG-IV show a significant decrease in cell viability with prior treatment of HAMI3379 followed by EC50’s of Bleomycin, Panobinostat and Vincristine (** <0.01 and ***p=0.0001).

## Discussion

Currently the standard of care for children with pDMG; radiotherapy, is considered palliative. There are no effective treatments for children with pDMG. Until fairly recently, paediatric high-grade gliomas have been grouped with their adult counterparts, but due to the increase in stereotactic biopsies to obtain tissue for genomic sequencing it has become clear they are different (13,28,29). It has been suggested that pDMG’s may arise from an aborted cell differentiation program of the developing pons, resulting in uncontrolled proliferation, with the cell of origin still not known (2,8,30).

Brain development is a precise, regulated molecular, cellular and epigenetic process that is governed by a genetic blueprint and environmental factors. The main steps in brain development are proliferation of cells, migration, differentiation, myelination and the formation of synapses. When these processes are interrupted or malfunction and pathologies arise (9,31,32).

GPR17 is a potent targetable receptor that has been shown to promote oligodendrocyte differentiation and can be targeted in pDMG cells using an experimental antagonist, HAMI3379 (12). MDL 299,51 was first identified as a small molecule agonist by Hennen et al in a comprehensive *in vitro* study in human and rodent primary cells (31). The use of HAMI3379 as a small molecule antagonist was identified by the same group from the University of Bonn, with a similar methodology and employing MDL 299,51 as a GPR17 activator (32). Treatment with HAMI3379 causes a reduction in GPR17 expression, mRNA and protein, and resulting in an increase in O4. In addition, this treatment led to a reduction in sphere size, cell motility, cell velocity and self-renewal capacity. When pDMG cells had pre-treatment with HAMI3379 in combination with known cytotoxic agents, there was a decrease in cell viability compare to cytotoxic alone.

### GPR17 as a therapeutic target for DIPG and clinical implications

The targeting of GPR17 using a small molecule antagonist has shown promise in modulating pDMG genotype and phenotype *in vitro.* The use of HAMI3379 alone does not cause a significant effect on cell viability in pDMG cells, but in combination with potential therapies it does. HAMI3379 is an experimental drug that needs to undergo further testing before it can be used in the clinic but shows promise to target GPR17 in pDMG. There has been no change in DIPG patient survival in over 50 years and patients do not benefit from systemic chemotherapy. This is thought to be due to a combination of the anatomical location, an intact blood brain barrier and inadequate drug delivery systems (33,34). The delivery of chemotherapeutics systemically (intravenously or orally) fail to reach adequate concentrations in the tumour. Development of local administration techniques using convection-enhanced delivery (CED) to deliver drugs directly into the tumour, bypassing the blood brain barrier are promising techniques (35). CED is an attractive form of delivery as it ensures the therapeutic reaches the tumour, hopefully limiting the effect on the normal brain. CED currently undergoing a number of clinical trials in brain tumours and Parkinson’s disease (33,36,37).

Cancers have an ability to adapt to changes in their environment to be supportive of proliferation and survival. Paediatric brain cancers have the added complexity of the brain undergoing dynamic changes in normal development, these need to be considered in the development of therapies. A combination of treatments that specifically target pDMG cells in the precious environment of the developing brain, rendering them more susceptible to cytotoxics agents may provide an improved treatment strategy.

By using real patient data, we identified GPR17 to be dysregulated in pDMG patients and confirmed this in our cell lines and interrogate its role further by the utilisation of a small molecule antagonist and agonist to modulate the expression and activity of GPR17.

As this work was being finalised for submission, a comprehensive and systematic computational investigation using RNA-Seq analysis of 169 glioblastoma (GBM) samples with a focus on GPR17 was published (38). In addition to showing the differential expression of GPR17 and over 150 genes in GBM, the authors provide data to support the role of GPR17 in multiple tumour promoting signalling pathways.

## Materials and methods

### Bioinformatics

Patient data and gene expression datasets were obtained from R2: microarray analysis and visualization platform (http://hgserver1.amc.nl/cgi-bin/r2/main.cgi). Datasets used in this study were Paugh (GSE26576) (39) and Pfister (GSE36245) (40).

### Cell culture

The VUMC-DIPG-A and VUMC-DIPG-08 cell lines were kindly provided by Dr.Ester Hulleman (VUmc Cancer Center, Amsterdam, The Netherlands). SU-DIPG-IV cells were kindly provided by Dr. Michelle Monje (Stanford University, California, USA). The use of these cells at the University of Portsmouth is possible through an arranged Material Transfer Agreement (MTA). DIPG cell lines were cultured in serum free conditions in medium designated “Tumour Stem Media (TSM)”, consisting of DMEM/F12, Neurobasal(-A) (Invitrogen), B27(-A) (Invitrogen), human-bFGF (20 ng/mL), human-EGF (20 ng/mL) (Miltenyi), human PDGF-AA (10 ng/mL) and PDGF-BB (10 ng/mL) (Peprotech) and heparin (2 ng/mL) (Sigma Aldrich).

### RT-qPCR

RNA was extracted using RNeasy Mini Kit (Qiagen) according to manufacturer’s instructions. Twenty-five nanograms of total RNA was used per reaction with PrecisionPLUS OneStep RT-qPCR Master Mix (PrimerDesign, UK) according to manufacturer’s instructions. Samples were run using a Roche Lightcycler 96 (Roche) instrument according to manufacturer’s instructions. Primers are as follows: *GPR17 (*fwd 5-CTCTGACTCCAGCCAAAGCATG-3, rev 5-GGTAGAAGGAGGCGAACAGCAT-3) and GAPDH (fwd 5-GTCTCCTCTGACTTCAACAGCG-3, rev 5-ACCACCCTGTTGCTGTAGCCAA-3).Data normalisation was performed using the housekeeping gene GAPDH and further analysis was carried out using the 2−ΔΔCT method (41).

### Flow cytometry

Cells were plated at 2 × 10^3^/ well in 6-well plates overnight, then treated with MDL 299,51 or HAMI3379 for 48 hours. Cells were collected via gentle scraping, blocked in 2% goat serum for 1 hour at room temperature. For intracellular antigens, cells were fixed and permeabilised with Cytofix/Cytoperm (BD Biosciences) for 20 minutes at 4°C. Fixed cells were incubated in primary antibodies for 1 hour at room temperature, washed thoroughly, then incubated with fluorochrome-conjugated secondary antibodies (1:500, Life Technologies) for 30 minutes at room temperature. Samples were analysed on BD FACSCalibur™, using CellQuestPro software. Each experiment was repeated three independent times in triplicate. Data were represented as percentage of positive cell population and mean fluorescence intensity (MFI).

### Immunocytochemistry

Cells were plated on to sterile glass cover slips at 1 × 10^3^/ well in 6 well plates overnight and then treated with MDL 299,51 or HAMI3379 for 48 hours. Cells were fixed with 4% paraformaldehyde (PFA) (Sigma, UK). For intracellular antigens, cells were permeabilised with 0.01% Triton-X. Non-specific antigens were blocked with 10% goat serum (Bio-Sera) for 1 hour at room temperature. Fixed cells were incubated in primary antibodies overnight at 4°C, then washed thoroughly with PBS then incubated with fluorochrome-conjugated AlexaFluor^®^ secondary antibodies (1:500, Life Technologies) for 1 hour at room temperature. Hoechst Blue (Sigma, UK) was used as nuclear counterstain. Coverslips were mounted on slides with Vectashield mounting medium (Vector Labs). Fluorescent images were obtained using Zeiss Axio Imager ZI fluorescent microscope using Volocity 6.1 acquisition software (PerkinElmer) or Zeiss LSM 510 Meta Axioskop2 confocal microscope and Zen software (Zeiss).

### MTS

Cells were plated at 1 × 10^3^/well overnight in a 96 well plate, then treated with either MDL 299,51 or HAMI3379. Ten microlitres of Cell Titre 96^®^ Aqueous One Solution was added to each well. The plate was incubated at 37°C for 2 hours before the absorbance is measured at 490nm using a POLARstar OPTIMA plate reader (BMG Labtech, UK). Readings were blank corrected by subtracting cell free controls and data normalised to untreated control.

### Live cell imaging

Cells were plated at 2 × 10^3^/ well in 6-well plates overnight, then treated with MDL 299,51 or HAMI3379 and live cell imaging was performed using a Zeiss Axiovert 200M (inverted) microscope (Carl Zeiss, Welwyn Garden City, Herts, UK) contained in an incubator (37 °C, 5% CO2 / 21% O2, humid atmosphere). A 5x objective was used and phase images were acquired every 30 min for 72 h (V5.4, Perkin Elmer). Single cells were manual tracked using Volocity software (V6.1.1, Perkin Elmer) for the course of the experiment, at least 10 cells per well were tracked.

### Statistical analysis

All experiments were performed at least three times in triplicate, and data expressed as mean ± standard error of mean (SEM). All statistical analyses were performed using Graph Pad Prism 6 software.

## Acknowledgments

We would like to sincerely thank Amanda at Abbie’s Army, Cassandra at Cure4cam, Harry St Ledger and his family and friends, and special thank you to Sarah and Barbara from Ollie Young Foundation. You all inspire us and together we will make a difference. Special thanks to Professor Geoffrey Pilkington for his support of this project and the DIPG cell lines provided by Dr.Ester Hulleman and Dr. Michelle Monje.

## Funding

This work was supported by funding from the following charities: Ollie Young Foundation and the University of Portsmouth (KFL), Brain Tumour Research (HLF), Abbie’s Army (HLF), Cure4cam (HLF).

## Notes

**Conflicts of Interest** The authors declare no potential conflicts of interest.

### Competing Interest Statement

The authors have declared no competing interest.

## References

1. Ballester LY, Wang Z, Shandilya S, Miettinen M, Burger PC, Eberhart CG, et al. Morphologic Characteristics and Immunohistochemical Profile of Diffuse Intrinsic Pontine Gliomas. Am J Surg Pathol [Internet]. 2013;37:1357–64. Available from: http://content.wkhealth.com/linkback/openurl?sid=WKPTLP:landingpage&an=00000478-201309000-00008

2. Baker SJ, Ellison DW, Gutmann DH. Pediatric gliomas as neurodevelopmental disorders. Glia [Internet]. 2015/12/08. 2016;64:879–95. Available from: http://onlinelibrary.wiley.com/store/10.1002/glia.22945/asset/glia22945.pdf?v=1&t=iuzl7na4&s=b0550881fa5fa2bc825c3bc9ddd07f327f7b6a0a

3. Mackay A, Burford A, Carvalho D, Izquierdo E, Fazal-Salom J, Taylor KR, et al. Integrated Molecular Meta-Analysis of 1,000 Pediatric High-Grade and Diffuse Intrinsic Pontine Glioma. Cancer Cell. 2017;32:520–537.e5.

4. Sengupta S, Sobo M, Lee K, Senthil Kumar S, White AR, Mender I, et al. Induced Telomere Damage to Treat Telomerase Expressing Therapy-Resistant Pediatric Brain Tumors. Mol Cancer Ther [Internet]. 2018 [cited 2018 May 5];molcanther.0792.2017. Available from: http://www.ncbi.nlm.nih.gov/pubmed/29654065

5. Ozawa PM, Ariza CB, Ishibashi CM, Fujita TC, Banin-Hirata BK, Oda JM, et al. Role of CXCL12 and CXCR4 in normal cerebellar development and medulloblastoma. Int J Cancer [Internet]. 2016;138:10–3. Available from: http://www.ncbi.nlm.nih.gov/pubmed/25400097

6. Wang L, Li Z, Zhang M, Piao Y, Chen L, Liang H, et al. H3 K27M-mutant diffuse midline gliomas in different anatomical locations. Hum Pathol. United States; 2018;78:89–96.

7. Louis DN, Perry A, Reifenberger G, von Deimling A, Figarella-Branger D, Cavenee WK, et al. The 2016 World Health Organization Classification of Tumors of the Central Nervous System: a summary. Acta Neuropathol [Internet]. 2016;131:803–20. Available from: http://www.ncbi.nlm.nih.gov/pubmed/27157931

8. Jones C, Karajannis MA, Jones DT, Kieran MW, Monje M, Baker SJ, et al. Pediatric high-grade glioma: biologically and clinically in need of new thinking. Neuro Oncol [Internet]. 2016; Available from: http://www.ncbi.nlm.nih.gov/pubmed/27282398

9. Tate MC, Lindquist RA, Nguyen T, Sanai N, Barkovich AJ, Huang EJ, et al. Postnatal growth of the human pons: a morphometric and immunohistochemical analysis. J Comp Neurol [Internet]. 2015;523:449–62. Available from: http://www.ncbi.nlm.nih.gov/pubmed/25307966

10. Fields RD. A new mechanism of nervous system plasticity: Activity-dependent myelination. Nat. Rev. Neurosci. 2015. page 756–67.

11. Mount CW, Monje M. Wrapped to Adapt: Experience-Dependent Myelination. Neuron. 2017. page 743–56.

12. Merten N, Fischer J, Simon K, Zhang L, Schröder R, Peters L, et al. Repurposing HAMI3379 to Block GPR17 and Promote Rodent and Human Oligodendrocyte Differentiation. Cell Chem Biol. 2018;

13. Schroeder KM, Hoeman CM, Becher OJ. Children are not just little adults: recent advances in understanding of diffuse intrinsic pontine glioma biology. Pediatr Res [Internet]. International Pediatric Research Foundation, Inc.; 2014;75:205–9. Available from: http://dx.doi.org/10.1038/pr.2013.194

14. Marucci G, Dal Ben D, Lambertucci C, Santinelli C, Spinaci A, Thomas A, et al. The G Protein-Coupled Receptor GPR17: Overview and Update. ChemMedChem [Internet]. 2016;11:2567–74. Available from: http://www.ncbi.nlm.nih.gov/pubmed/27863043

15. Fumagalli M, Daniele S, Lecca D, Lee PR, Parravicini C, Douglas Fields R, et al. Phenotypic changes, signaling pathway, and functional correlates of GPR17-expressing neural precursor cells during oligodendrocyte differentiation. J Biol Chem. 2011;286:10593–604.

16. Ciana P, Fumagalli M, Trincavelli ML, Verderio C, Rosa P, Lecca D, et al. The orphan receptor GPR17 identified as a new dual uracil nucleotides/cysteinyl-leukotrienes receptor. EMBO J [Internet]. 2006;25:4615–27. Available from: http://emboj.embopress.org/cgi/doi/10.1038/sj.emboj.7601341

17. Lecca D, Trincavelli ML, Gelosa P, Sironi L, Ciana P, Fumagalli M, et al. The recently identified P2Y-like receptor GPR17 is a sensor of brain damage and a new target for brain repair. PLoS One [Internet]. 2008;3:e3579. Available from: http://www.ncbi.nlm.nih.gov/pubmed/18974869

18. Boda E, Hoxha E, Montarolo F, Rosa P, Abbracchio MP, Buffo A, et al. The P2Y-like GPR17 receptor participates in oligodendrocyte precursor cell reaction in a model of chronic cerebral amyloidosis. Alzheimer’s Dement. 2009. page P490.

19. Chen Y, Wu H, Wang S, Koito H, Li J, Ye F, et al. The oligodendrocyte-specific G protein-coupled receptor GPR17 is a cell-intrinsic timer of myelination. Nat Neurosci [Internet]. 2009;12:1398–406. Available from: http://www.ncbi.nlm.nih.gov/pubmed/19838178

20. Khan MZ, He L. Neuro-psychopharmacological perspective of Orphan receptors of Rhodopsin (class A) family of G protein-coupled receptors. Psychopharmacol [Internet]. 2017;234:1181–207. Available from: http://www.ncbi.nlm.nih.gov/pubmed/28289782

21. Shi QJ, Wang H, Liu ZX, Fang SH, Song XM, Lu YB, et al. HAMI 3379, a CysLT 2 R antagonist, dose- and time-dependently attenuates brain injury and inhibits microglial inflammation after focal cerebral ischemia in rats. Neuroscience. 2015;

22. Fumagalli M, Daniele S, Lecca D, Lee PR, Parravicini C, Fields RD, et al. Phenotypic changes, signaling pathway, and functional correlates of GPR17-expressing neural precursor cells during oligodendrocyte differentiation. J Biol Chem [Internet]. 2011;286:10593–604. Available from: http://www.ncbi.nlm.nih.gov/pubmed/21209081

23. La Manno G, Gyllborg D, Codeluppi S, Nishimura K, Salto C, Zeisel A, et al. Molecular Diversity of Midbrain Development in Mouse, Human, and Stem Cells. Cell. 2016;167:566–580.e19.

24. Kang HJ, Kawasawa YI, Cheng F, Zhu Y, Xu X, Li M, et al. Spatio-temporal transcriptome of the human brain. Nature. 2011;

25. Paugh BS, Zhu X, Qu C, Endersby R, Diaz AK, Zhang J, et al. Novel oncogenic PDGFRA mutations in pediatric high-grade gliomas. Cancer Res. 2013;

26. Filbin MG, Tirosh I, Hovestadt V, Shaw ML, Escalante LE, Mathewson ND, et al. Developmental and oncogenic programs in H3K27M gliomas dissected by single-cell RNA-seq. Science (80-). 2018;

27. Hennen S, Wang H, Peters L, Merten N, Simon K, Spinrath A, et al. Decoding Signaling and Function of the Orphan G Protein-Coupled Receptor GPR17 with a Small-Molecule Agonist. Sci Signal [Internet]. 2013;6:ra93–ra93. Available from: http://stke.sciencemag.org/cgi/doi/10.1126/scisignal.2004350

28. Bjerke L, Mackay A, Nandhabalan M, Burford A, Jury A, Popov S, et al. Histone H3.3 mutations drive pediatric glioblastoma through upregulation of MYCN. Cancer Discov. 2013;3:512–9.

29. Khuong-Quang DA, Buczkowicz P, Rakopoulos P, Liu XY, Fontebasso AM, Bouffet E, et al. K27M mutation in histone H3.3 defines clinically and biologically distinct subgroups of pediatric diffuse intrinsic pontine gliomas. Acta Neuropathol. 2012;124:439–47.

30. Filbin MG, Tirosh I, Hovestadt V, Shaw ML, Escalante LE, Mathewson ND, et al. Developmental and oncogenic programs in H3K27M gliomas dissected by single-cell RNA-seq. Science [Internet]. American Association for the Advancement of Science; 2018 [cited 2018 Apr 25];360:331–5. Available from: http://www.ncbi.nlm.nih.gov/pubmed/29674595

31. Ouyang M, Dubois J, Yu Q, Mukherjee P, Huang H. Delineation of early brain development from fetuses to infants with diffusion MRI and beyond. Neuroimage [Internet]. Academic Press; 2018 [cited 2018 May 4]; Available from: https://www.sciencedirect.com/science/article/pii/S105381191830301X#fig1

32. Lindquist RA, Guinto CD, Rodas-Rodriguez JL, Fuentealba LC, Tate MC, Rowitch DH, et al. Identification of proliferative progenitors associated with prominent postnatal growth of the pons. Nat Commun. England; 2016;7:11628.

33. El-Khouly FE, van Vuurden DG, Stroink T, Hulleman E, Kaspers GJL, Hendrikse NH, et al. Effective drug delivery in diffuse intrinsic pontine glioma: A theoretical model to identify potential candidates. Front Oncol. 2017;7.

34. Warren KE. Beyond the Blood:Brain Barrier: The Importance of Central Nervous System (CNS) Pharmacokinetics for the Treatment of CNS Tumors, Including Diffuse Intrinsic Pontine Glioma. Front Oncol [Internet]. Switzerland; 2018;8:239. Available from: https://www.frontiersin.org/article/10.3389/fonc.2018.00239/full

35. Himes BT, Zhang L, Daniels DJ. Treatment strategies in diffuse midline gliomas with the H3K27M mutation: The role of convection-enhanced delivery in overcoming anatomic challenges. Front. Oncol. 2019.

36. Souweidane MM, Kramer K, Pandit-Taskar N, Zhou Z, Haque S, Zanzonico P, et al. Convection-enhanced delivery for diffuse intrinsic pontine glioma: a single-centre, dose-escalation, phase 1 trial. Lancet Oncol. 2018;

37. Whone A, Luz M, Boca M, Woolley M, Mooney L, Dharia S, et al. Randomized trial of intermittent intraputamenal glial cell line-derived neurotrophic factor in Parkinson’s disease. Brain. 2019;

38. Mutharasu G, Murugesan A, Konda Mani S, Yli-Harja O, Kandhavelu M. Transcriptomic analysis of glioblastoma multiforme providing new insights into GPR17 signaling communication. J Biomol Struct Dyn. England; 2020;1–14.

39. Paugh BS, Broniscer A, Qu C, Miller CP, Zhang J, Tatevossian RG, et al. Genome-wide analyses identify recurrent amplifications of receptor tyrosine kinases and cell-cycle regulatory genes in diffuse intrinsic pontine glioma. J Clin Oncol. 2011;29:3999–4006.

40. Sturm D, Witt H, Hovestadt V, Khuong-Quang DA, Jones DTW, Konermann C, et al. Hotspot Mutations in H3F3A and IDH1 Define Distinct Epigenetic and Biological Subgroups of Glioblastoma. Cancer Cell. 2012;

41. Livak KJ, Schmittgen TD. Analysis of relative gene expression data using real-time quantitative PCR and the 2-ΔΔCT method. Methods. 2001;

