## Supplemental data for "Targeting G protein-coupled receptor-17 (GPR17) upregulation in paediatric diffuse midbrain gliomas leads to altered phenotype and susceptibility to therapies"

<u>**Supplementary Data – Loveson and Fillmore 2020**</u>

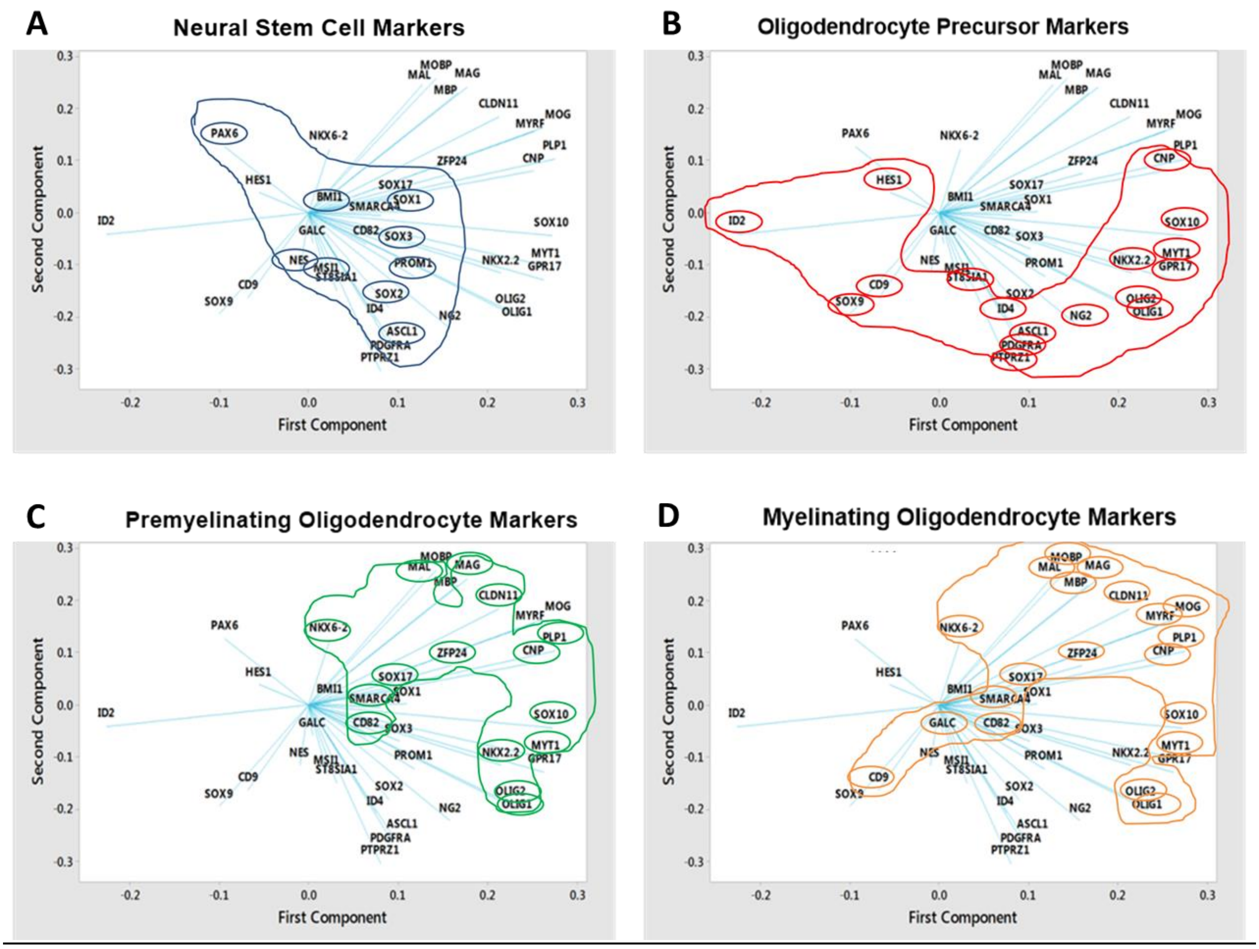


*Figure S 1 - Loading plot from PCA analysis with stage-specific genes highlighted. A) Neural stem cell markers B) Oligodendrocyte Precursor Markers C) Premyelinating Oligodendrocyte Markers and D) Myelinating Oligodendrocyte Markers.*

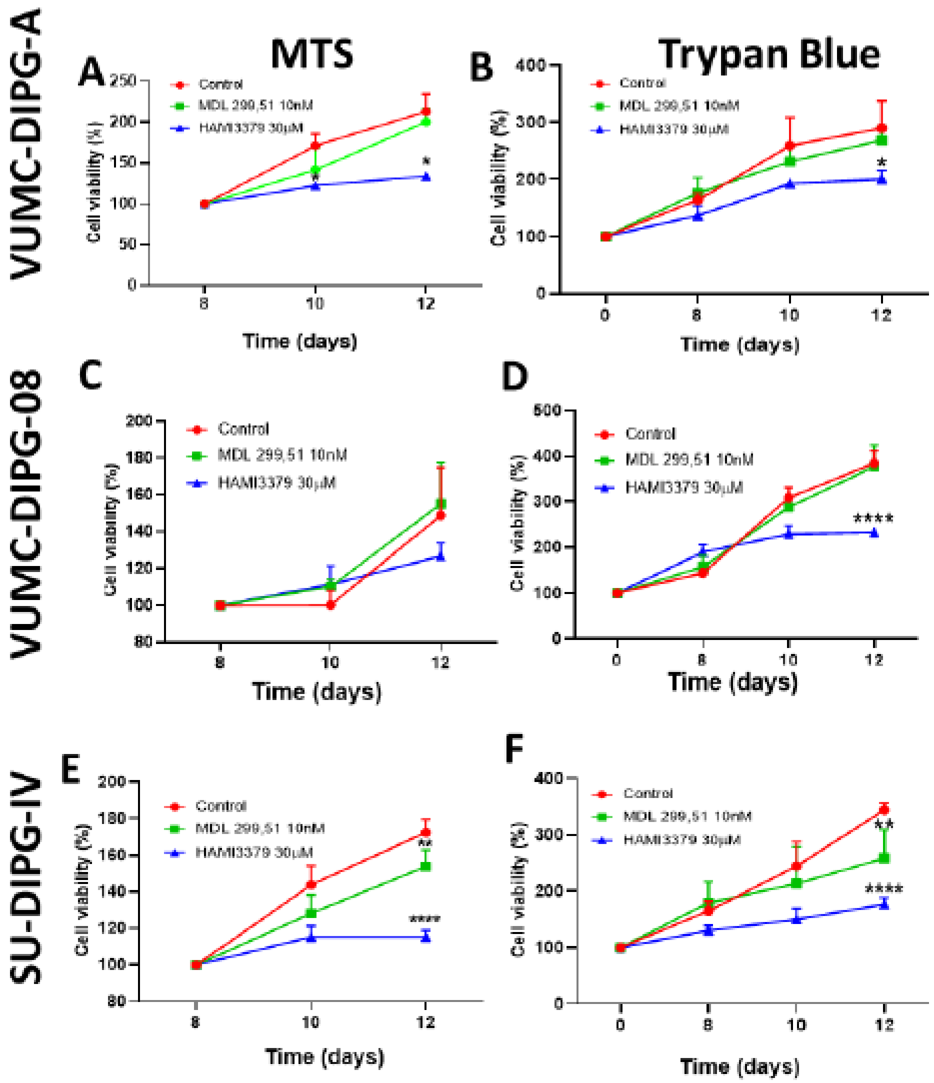


*Figure S 2 - Measurement of self-renewal capacity of DIPG cells. HAMI3379 reduces the self-renewal capacity of DIPG-cells. DIPG-cells were plated after 7 days of treatment with 10nM MDL299,51 or HAMI3379 and cell viability assays were performed. MTS assay performed on days 8, 10 and 12 after treatment and absorbance measured at 490nm (A, C, and E). Measurements were blank corrected and normalised to 8-day time. Trypan blue exclusion assay performed after 7 days of treatment and on days 8, 10 and 12 (B, D, and F). Cell number was normalised to day 7. Graphs show mean with SEM. Statistics were performed using GraphPad Prism, Two-way ANOVA with Dunnett's multiple comparison test. *p<0.04. ****p<0.0001.*

*Table S1-EC50's determined by sigmoidal dose-response*

| | Bleomycin (mg/mL) | Vincristine (nM) | Panobinostat (nM) |
|---|---|---|---|
| VUMC-DIPG-A | 1.87 | 15.29 | 48.97 |
| VUMC-DIPG-08 | 1.53 | 14.19 | 53.06 |
| SU-DIPG-IV | 1.38 | 11.85 | 31.94 |

*Table S2- STR Profiles of cell lines used in this study*

| | AMEL | CSF1PO | D13S317 | D16S539 | D21S11 | D5S818 | D7S820 | TH01 | TPOX | vWA |
|---|---|---|---|---|---|---|---|---|---|---|
| OPC | X, X | 11, 11 | 9,9 | 12, 13 | 30, 30 | 11, 12 | 9, 11 | 9.3, 9.3 | 10, 11 | 17, 17 |
| VUMC-DIPG-A | X, X | 12, 12 | 11, 11 | 8, 12 | 29, 29 | 11, 12 | 11, 11 | 6, 8 | 11, 11 | 16,17 |
| VUMC-DIPG-08 | X, Y | 9, 13 | 12, 12 | 11, 13 | 30, 30.2 | 12, 13 | 11, 13 | 6, 9.3 | 11, 12 | 16, 18 |
| SU-DIPG-IV | X, X | 9, 10 | 7, 12 | 9, 12 | 29, 31 | 12, 13 | 10, 11 | 6, 9.3 | 8, 8 | 15,19 |
